# Saccades Influence Functional Modularity in the Human Cortical Vision Network

**DOI:** 10.1101/2024.05.17.594722

**Authors:** G. Tomou, B. Baltaretu, A. Ghaderi, J. D. Crawford

**Author notes:** Corresponding Author: Professor J.D. Crawford, Centre for Vision Research, Room 0009A, Lassonde Bldg. York University, Toronto, Ontario, Canada, M3J 1P3.

## Abstract

Visual cortex is thought to show both dorsoventral and hemispheric modularity, but it is not known if the same functional modules emerge spontaneously from an unsupervised network analysis, or how they interact when saccades necessitate increased sharing of spatial information. Here, we address these issues by applying graph theory analysis to fMRI data obtained while human participants decided whether an object’s shape or orientation changed, with or without an intervening saccade across the object. BOLD activation from 50 vision-related cortical nodes was used to identify local and global network properties. Modularity analysis revealed three sub-networks during fixation: a bilateral parietofrontal network linking areas implicated in visuospatial processing and two lateralized occipitotemporal networks linking areas implicated in object feature processing. When horizontal saccades required visual comparisons between visual hemifields, functional interconnectivity and information transfer increased, and the two lateralized ventral modules became functionally integrated into a single bilateral sub-network. This network included ‘between module’ connectivity hubs in lateral intraparietal cortex and dorsomedial occipital areas previously implicated in transsaccadic integration. These results provide support for functional modularity in the visual system and show that the hemispheric sub-networks are modified and functionally integrated during saccades.

## INTRODUCTION

Vision supports various perceptual, cognitive, and sensorimotor processes by influencing signals throughout the cerebral cortex. Given the complexity and broad distribution of these signals, the visual system is often described in terms of *functional modularity*, i.e., divisions based on *hemispheric lateralization* of the visual fields (Sperry, 1961; Strother et al., 2017) and *dorsoventral streams* for fundamentally different computations (Goodale & Milner, 1992; Ungerleider & Mishkin, 1982). These concepts are largely based on traditional region-based, neurophysiological, neurpsychological, and neuroimaging approaches. This begs two questions: 1) does the cortical distribution of visual signals show the same modularity, and how are these signals communicated in tasks that require the integration of multiple modules? For example, in the presence of saccadic eye movements, the visual system is thought to engage both the dorsal and ventral streams (Prime et al., 2008) and often requires the remapping and sharing of information across visual fields (Medendorp et al., 2003; Merriam et al., 2007).

Anatomic and functional lateralization was one of the first properties discovered in the visual system and is now widely assumed as fact, at least early in the system. Anatomic projections from the thalamus to early visual cortex promote a clear contralateral visual field specificity in occipital cortex (Sperry, 1961; Strother et al., 2017). However, visual-field specificity is less clear at higher levels of brain function. For example, parietal cortex shows a propensity to code contralateral visual field in neuroimaging studies (Medendorp et al., 2003; Sereno et al., 2001) but parietal field specificity is less clear in neurophysiological signals and visuomotor deficits (Mooshagian et al., 2018, 2022).

Dorsoventral modularity was originally based on neurophysiological recordings in primates: Ungerleider and Mishkin (1982) proposed that the feedforward processing of vision is divided into a ventral (occipital-temporal) ‘what’ stream versus a dorsal (occipital-parietal) ‘where’ stream for vision. Goodale and Milner later argued that the better distinction is ‘what’ versus ‘how’, where the ventral stream is used for perception and the dorsal stream is used for action (Goodale & Milner, 1992; Goodale & Westwood, 2004; Milner, 2017; Milner & Goodale, 2008). Subsequent neuroimaging studies have supported the division of object recognition in ventrolateral occipital and temporal cortex (Grill-Spector et al., 2001; Kanwisher et al., 1996) as opposed to saccade and reach areas in posterior parietal cortex (Hagler et al., 2007; Levy et al., 2007; Vesia & Crawford, 2012). Dorsal stream areas like the parietal eye fields are closely associated with prefrontal areas, including the frontal eye fields, supplementary eye fields, and dorsolateral prefrontal cortex. Collectively, the latter areas form both the cortical saccade network (Crawford et al., 2011; Gaymard et al., 1998; Lynch & Tian, 2006) and the ‘dorsal attention network’, exerting top-down influences on visual cortex (Astafiev et al., 2003; Farrant & Uddin, 2015; Fox et al., 2006).

These functional divisions beg the question of how they interact for real world behavior. A prime example is transsaccadic vision: humans make saccades several times per second (Rayner, 1978, 1998) requiring the visual system to retain, compare and/or integrate information between different visual fixations (Melcher & Colby, 2008; Prime et al., 2007; Tas et al. 2012; Edwards et al. 2018). This is thought to involve a network that includes saccade signals from the frontal and parietal eye fields (Prime et al., 2008, 2011), and low level feature processing in inferior parietal cortex and dorsomedial occipital cortex (Baltaretu et al., 2020, 2021; Dunkley et al., 2016). Further, when saccades reverse the visual field of an object, this requires comparisons of information between the two hemispheres (Malik et al., 2015; Medendorp et al., 2003; Merriam et al., 2003, 2007), presumably via the corpus callosum (Berman & Colby, 2009).

Importantly, nearly everything we know about the modularity of cortical saccade and visual systems is based on ‘region-of-interest’ analysis: much less is known about the dynamics and topology of these systems at the network level. Recently there has been increased focus on *functional ‘connectivity’*, i.e., signal correlations between different cortical regions referred to as nodes (Friston, 1994). Functional connectivity can be determined for specific ‘seed’ regions (e.g., Baltaretu et al., 2020, 2021, 2023), but more sophisticated techniques like graph theory analysis (GTA) can compute global network properties, including important network ‘hubs’ and formal measures of functional integration and segregation (Ghaderi et al., 2023; Rubinov & Sporns, 2010). Most important for our purposes, GTA measures of optimal community structure (groups of highly correlated regions that have few signal correlations with nodes outside of their community) allow one to determine modular organization of cortical regions into functional sub-networks (Boccaletti et al., 2006; Bullmore & Sporns, 2009; Newman, 2004; Rubinov & Sporns, 2010). Typically, these techniques have been applied to the visual system using ‘resting state’ imaging data (Reavis et al., 2020; Stevens et al., 2010), but we have recently shown that GTA can also be applied to sensory and motor events recorded during active behavior (Ghaderi et al., 2023).

Here, we performed a GTA / modularity analysis on functional magnetic resonance imaging (fMRI) data collected in a task that involved discriminating changes in the shape versus orientation of a central object with or without an intervening saccade that reversed the visual hemifield of the object (Fig 1a). A previous region of interest analysis of these data confirmed the role of dorsomedial occipital and inferior parietal cortex in transsaccadic feature integration (Baltaretu et al., 2023), Here, we derived fMRI time series data across a broad distribution of cortical ‘nodes’ in the visual system (Baltaretu et al., 2020; L. Wang et al., 2015), and then used GTA to derive formal measures of functional modularity, integration, and segregation between these nodes. With these techniques, we investigated two questions: 1) will lateralization and dorsal-ventral modularity automatically arise from this task and analysis, 2) will saccades increase functional connectivity in the visual system, specifically across hemispheres.

## RESULTS

Figure 1a shows our experimental paradigm. 17 participants fixated either left or right of center and were presented with an oriented object in their periphery. Following a brief mask, an object was once again presented; this object was either the same shape as the first presentation with a new orientation, or a new shape with the same orientation. During *Fixation* trials, gaze remained stable at the original location, but during *Saccade* trials, an eye movement was made to the opposite side, such that the second object appeared in the opposing visual hemifield. The latter condition thus required comparisons between visual information from the same location in space but viewed in oppositive hemifields. Experiments were performed within an MRI scanner in complete darkness (see Baltaretu et al., 2023, and *Materials and Methods* for details of experimental methodology). The data were used for a standard univariate / ROI analysis (Baltaretu et al., 2023) then were selected for this study because they provide a rich set of task-related signals for GTA.

**Figure 1.**
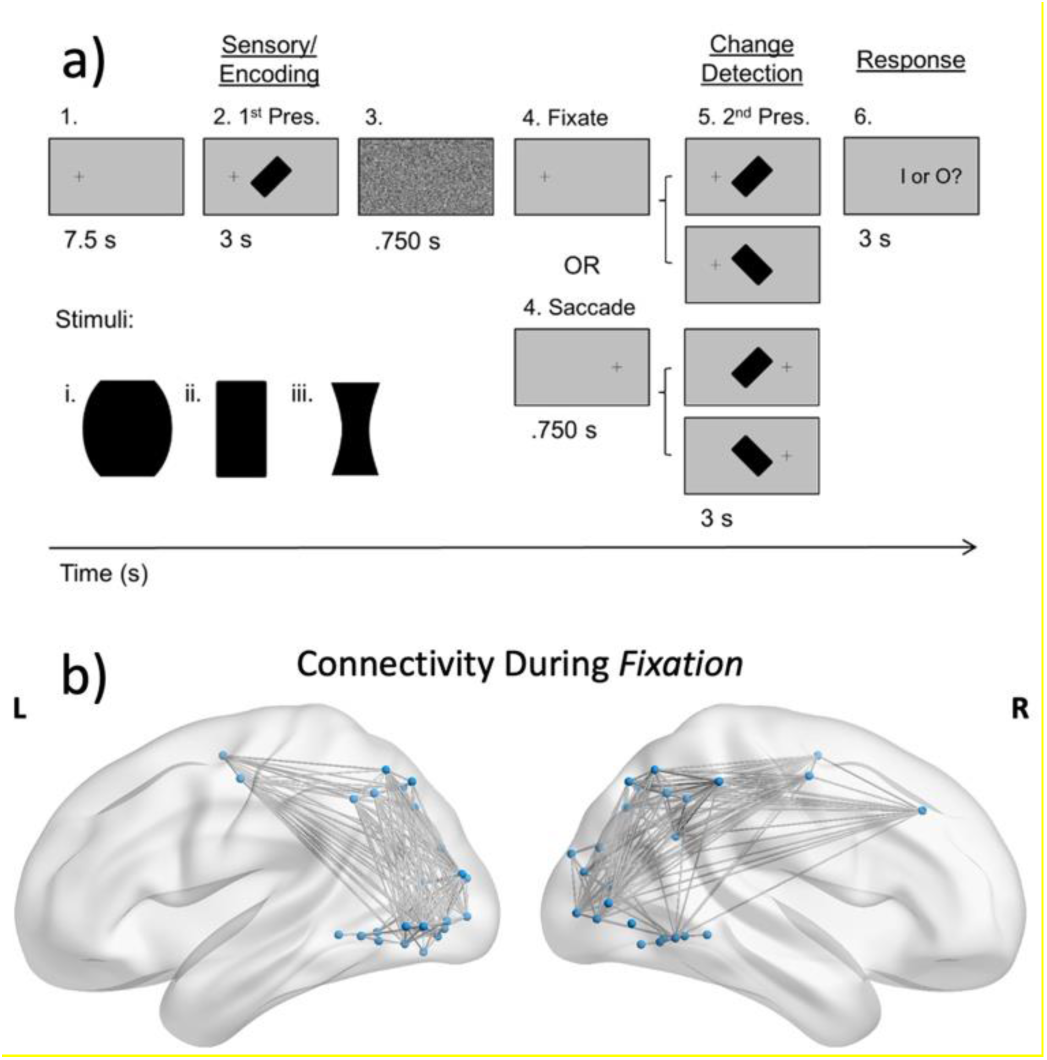
a) The event-related fMRI experiential paradigm that was used to identify cortical regions involved in object identity and orientation recognition and the influence of saccades on these processes. b) The 50 ROIs that were subjected to the GTA analysis and their correlated activity during *Fixation* trials. These regions are highlighted in *Table 1*.

To compute GTA measures, we derived group blood oxygenation level-dependent (BOLD) time series from 50 regions of interest (defined here as *nodes*), using coordinates derived from previous studies of the visual system (Fig 1b; Table 1). Superscript symbols^†^ indicate overlap with visual field-specific activation derived from the same dataset (Baltaretu et al., 2023). We used the *Newman modularity* approach which groups a network’s nodes into modules based on high degrees of correlated activation (Newman, 2004, 2006) to evaluate the degree of sub-networks within our network. Then we compared the resulting modules between our *Fixation* and *Saccade* conditions. To investigate the contribution of each node in information sharing across the network, we calculated their *Betweenness Centrality* and compared values between conditions. Higher centrality suggests greater involvement in neural information propagation through the network (Avena-Koenigsberger et al., 2017; Bullmore & Sporns, 2009). Finally, we calculated measures of functional integration (e.g., *global efficiency*) and segregation (e.g., *clustering coefficient*) to compare the properties of the overall network in our two conditions. It is important to note that none of these measures imply causality or anatomic connection, but rather patterns of signal correlation that can be interpreted relative to the known anatomy and functions of the system.

### Modularity and Eigenvector Centrality in the Fixation Task

Our first goal was to determine if functional connectivity in the *Fixation* task shows the kind of modularity suggested by traditional region-of-interest approaches. Figure 2 shows the results of an unsupervised Newman modularity analysis on our 50 nodes. This produced three sub-modules: a dorsal sub-network and two lateralized ventroposterior modules. The modules are shown as color coded ‘nodes’ (small dots), ‘edges’ (lines between the nodes), and hubs (large dots) based on *Eigenvector Centrality*, a measure of the importance of the node within its module.

The dorsal module showed bilateral connectivity that spanned our parietofrontal nodes in both cortical hemispheres. This module generally corresponded to the dorsal attention network (Astafiev et al., 2003; Farrant & Uddin, 2015; Fox et al., 2006), including nodes associated with visuospatial memory and eye movements such as right dorsolateral prefrontal cortex (Curtis & D’Esposito, 2003) the frontal eye fields (Prime et al., 2010, 2011), and several sites along the intraparietal sulci (Barash et al., 1991; Grefkes & Fink, 2005). Hubs determined by Eigenvector Centrality in this module included intraparietal sulcus (IPS1, IPS2, IPS3, and IPS4) and superior parietal lobule (SPL1).

The more posterior-ventral network was divided into two largely lateralized modules that encompassed our nodes from occipital and temporal cortex. The (mostly) left module included all nodes from the hemisphere but also some regions in the right hemisphere, whereas the other, smaller module fell exclusively within the right hemisphere. Overall, these modules included occipital / temporal nodes associated with low-to-high level object feature processing (Aminoff et al., 2013), but also some superior occipital nodes traditionally associated with the dorsal visual stream (Grill-Spector et al., 1999). The (mostly) leftward module included important Hubs in lateral occipital cortex and visual area V3A whereas the rightward module Hubs included visual areas V3A and V7/ISPO.

**Figure 2.**
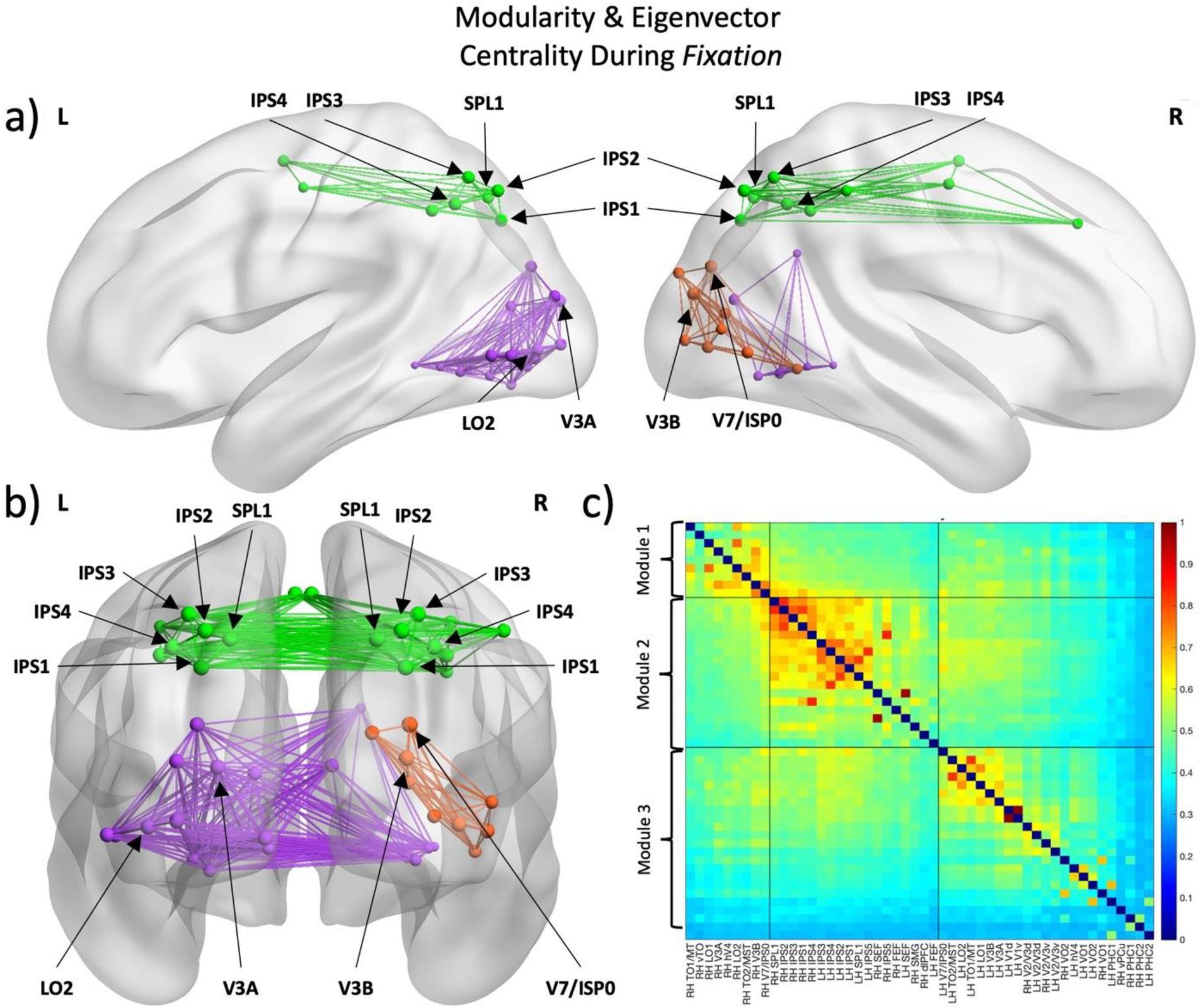
Modularity results of the *Fixation* condition. a) Lateral views highlight ventral-dorsal modularity in the visual system. Node size demonstrates Eigenvector Centrality with larger nodes being more central within each module (i.e., hub regions). Hubs were computed by normalizing eigenvector values and selecting those with values between 95-100. b) Posterior view highlights right-left lateralization modules. c) Adjacency matrices reorganized following modularity analysis. Black lines indicate separation of the ROIs into 3 modules, with the large black squares along the diagonal showing distinct modular groups.

### Modularity and Eigenvector Centrality in the Saccade Task

Our next goal was to determine the influence of saccades (across the visual object) on the modular networks described in the previous section. Figure 3 shows the results of a Newman modularity analysis of the fMRI data from our saccade condition. Overall, dorso-ventral modularity was preserved during saccades (Fig 3), except two regions (bilateral V7/IPS0) included in the dorsal fixation module that became more correlated with the ventral modules during saccades. In general, hubs were similar in the saccade and fixation condition, but we will quantify this more carefully below.

Perhaps more importantly, during saccades the two ventral modules combined into a single bilateral module. This is most clear from the posterior view of the brain (Fig 3b). Recall that in our paradigm, saccades reversed the visual hemifield (and thus the cortical hemisphere) used to view the second visual stimulus relative to the first stimulus, before participants had to compare these stimuli. Figure 3 suggests this leads to increased communication, or sharing of similar information, between right and left visual cortex.

**Figure 3.**
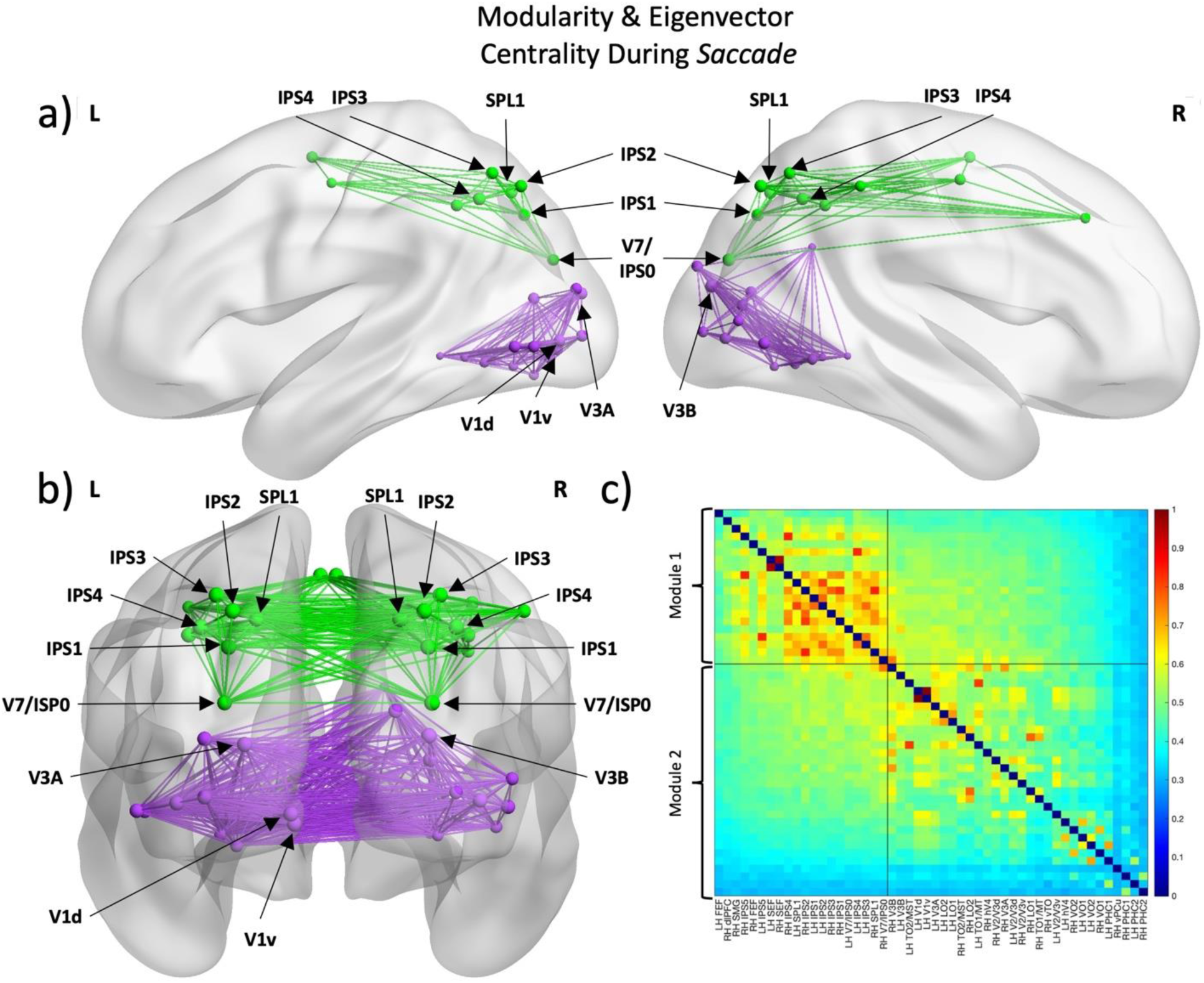
Modularity results of the *Saccade* condition. a) Lateral views highlight ventral-dorsal modularity in the visual system. Node size demonstrates Eigenvector Centrality with larger nodes being more central within each module (i.e., hub regions). Hubs were computed by normalizing eigenvector values and selecting those with values between 95-100. b) Posterior view indicates that the two right-left lateralized modules become highly intercorrelated, resulting in a single module. c) Adjacency matrices reorganized following modularity analysis. Black lines indicate separation of the ROIs into 2 modules, with the large black squares along the diagonal showing distinct modular groups.

### Communication Between Modules: Betweenness Centrality

The application of Eigenvector Centrality to cortical nodes in in Figures 2 and 3 inform one about the importance of certain hubs *within* modules. Assuming these modules are not completely independent, it was also of interest to determine which nodes of each module act as important hub regions that ‘bridge’ *between* modules. *Betweenness Centrality* is a measure that determines which nodes facilitate functional integration between modules (Avena-Koenigsberger et al., 2017; Bullmore & Sporns, 2009). Figure 4 denotes the nodes with greatest Betweenness Centrality of each module in the *Fixation* and *Saccade* conditions, respectively. During fixation (Fig 4a), bilateral intraparietal sulcus areas IPS3 and IPS4 were identified as important Betweenness Centrality hubs for the bilateral dorsal (green) module, whereas bilateral V3B played this role in the two ventral networks. During saccades (Fig 4b), only bilateral intraparietal sulcus (IPS4) and right dorsomedial occipital cortex (V3B) were preserved as important hubs for communication between the two remaining dorsal and ventral modules. Both posterior parietal and dorsomedial occipital cortex have been implicated in transsaccadic feature integration (Baltaretu et al., 2021, 2023; Dunkley et al., 2016; Prime et al., 2008; Schmitt et al., 2020).

**Figure 4.**
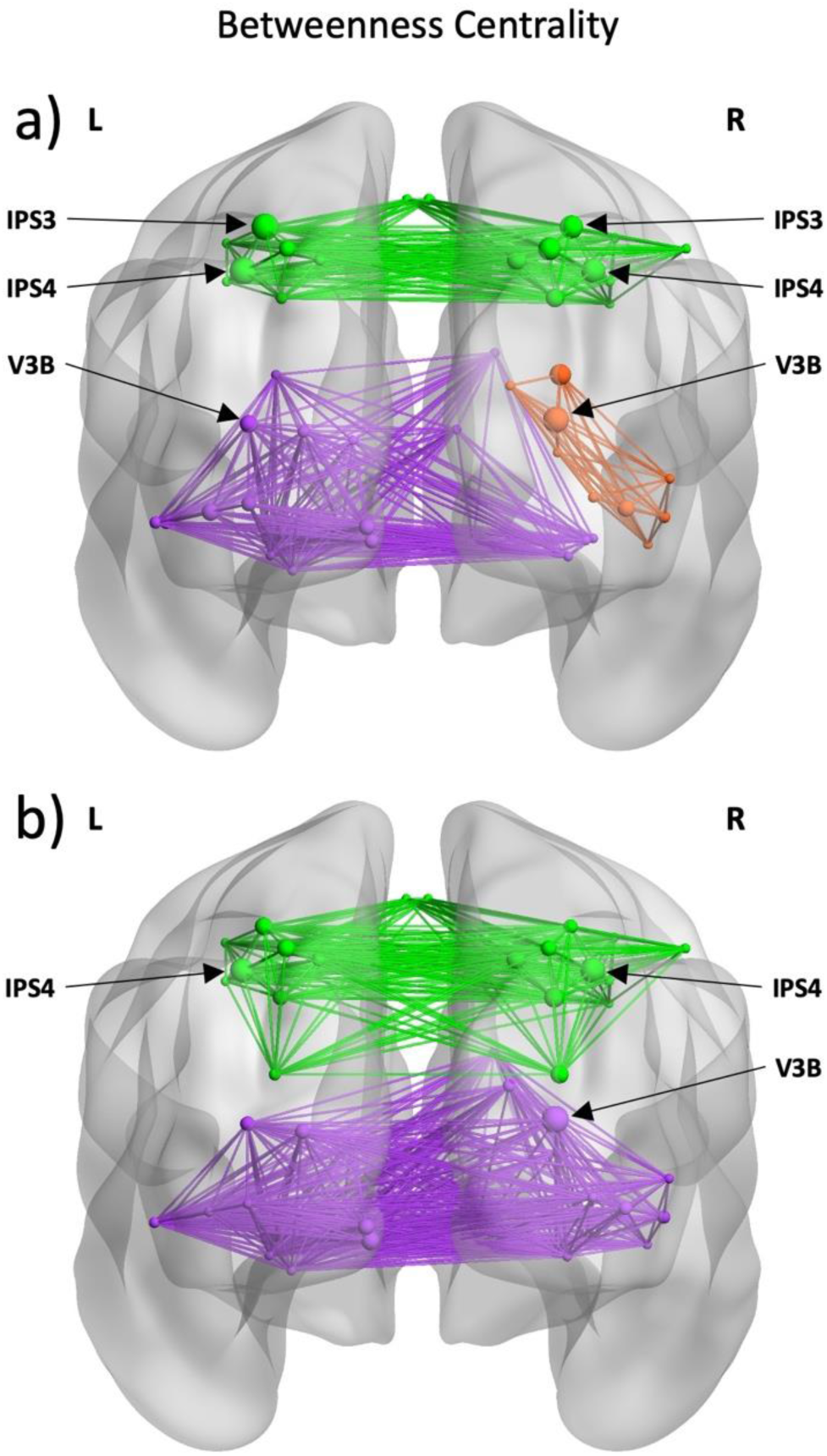
Betweenness Centrality hubs are labeled by normalizing betweenness values and selecting those with normalized values between 85-100. a) Modularity and Betweenness Centrality results of the *Fixation* condition. None of the nodes in the purple module satisfied the normalized betweenness value criteria; left V3B is labeled as the node with the highest Betweenness Centrality in the purple module. b) Modularity and Betweenness Centrality results of the *Saccade* condition.

### Fixation Versus Saccade Networks: Quantitative Comparison

In these final results we provide direct statistical comparisons between the fixation and saccade data described above to quantify the influence of saccades on network modularity, specific hubs, and global network properties.

**Eigenvector Centrality.** Figure 5 provides a direct comparison between Hubs in the Fixation and Saccade conditions (see Figure 5 caption for detailed statistics). We found that the saccade condition showed significantly higher Eigenvector Centrality in four of our occipital nodes, including 3 in the left hemisphere (V1v, V2/V3v, V1d, and V2/V3d) and one in the right hemisphere – V2/V3v. As noted above, both early visual cortex (Malik et al., 2015) and dorsomedial occipital cortex (Baltaretu et al., 2021, 2023) have previously been implicated in transsaccadic feature processing. Note that we performed a similar analysis on our Betweenness Centrality measures but found no significant difference.

**Figure 5.**
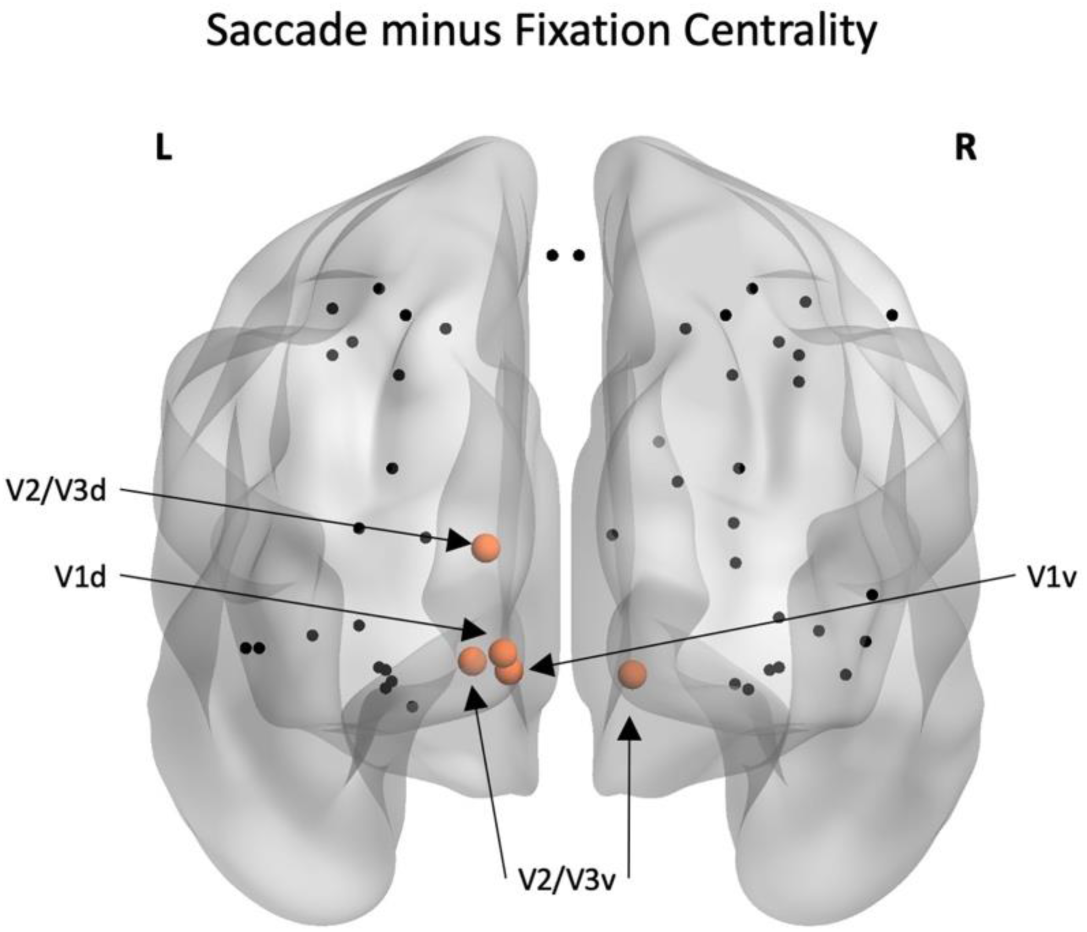
Node Eigenvector Centrality compared across the *Fixation* and *Saccade* conditions. The four regions in orange were significantly more central during *Saccade* relative to *Fixation*. Conditions were compared using nonparametric permutation t-tests and false detection rate (FDR) was used to control for multiple comparisons. Adjusted p-values (p_adj_ < 0.05) are reported here. These regions include 4 regions of the left hemisphere – V1v [saccade (M = 0.152, SD = 0.008); fixation (M = 0.148, SD = 0.009); t(16) = -0.007, p_adj_ = .008], V2/V3v [saccade (M = 0.137, SD = 0.008); fixation (M = 0.129, SD = 0.007); t(16) = -0.010, p_adj_ = .002], V1d [saccade (M = 0.153, SD = 0.009); fixation (M = 0.149, SD = 0.008); t(16) = -0.007, p_adj_ = .008], and V2/V3d [saccade (M = 0.143, SD = 0.006); fixation (M = 0.138, SD = 0.008); t(16) = -0.009, p_adj_ = .004] – and a single region of the right hemisphere – V2/V3v [saccade (M = 0.142, SD = 0.007); fixation (M = 0.135, SD = 0.008); t(16) = -0.010, p_adj_ = .002].

**Modularity**. To quantify the extent to which modularity was altered during saccades, we conducted a quantitative analysis on the sets of nodes that were identified to be part of the dorsal and ventral pathways during both conditions (excluding bilateral V7/IPS0, since these where the only nodes that changed their activity between both pathways). There was no significant change in the degree of modularity for the nodes in the dorsal pathway. However, in the ventral nodes showed increased modularity during saccades (*M* = 0.951, *SD* = 0.05) compared to fixation (*M* = 0.941, *SD* = 0.04), *t*(16) = -0.017, *p* = 0.002, (nonparametric t-test) in the ventral pathway, indicating significantly increased functional connectivity and information propagation among these regions (Fig 6a).

**Global Network Features.** In graph theory analysis, the *Cluster Coefficient* provides a measure of *functional segregation* (degree of interconnectivity at a local scale) whereas *Global Efficiency*, is a measure of functional integration (interconnectivity across the entire network); see Methods for details. Both these measures showed significant differences between our fixation and saccade data (Fig 6b). Τhere was a significant increase in *clustering coefficient* during saccades (*M* = 0.475, *SD* = 0.045) compared to fixation (*M* = 0.465, *SD* = 0.039), *t*(16) = -0.02, *p* = 0.036, and *global efficiency* during saccades (*M* = 0.481, *SD* = 0.045) compared to fixation (*M* = 0.471, *SD* = 0.038), *t*(16) = -0.02, *p* = 0.037, indicating that eye movements elicited both greater functional segregation and functional integration of the network respectively.

**Figure 6.**
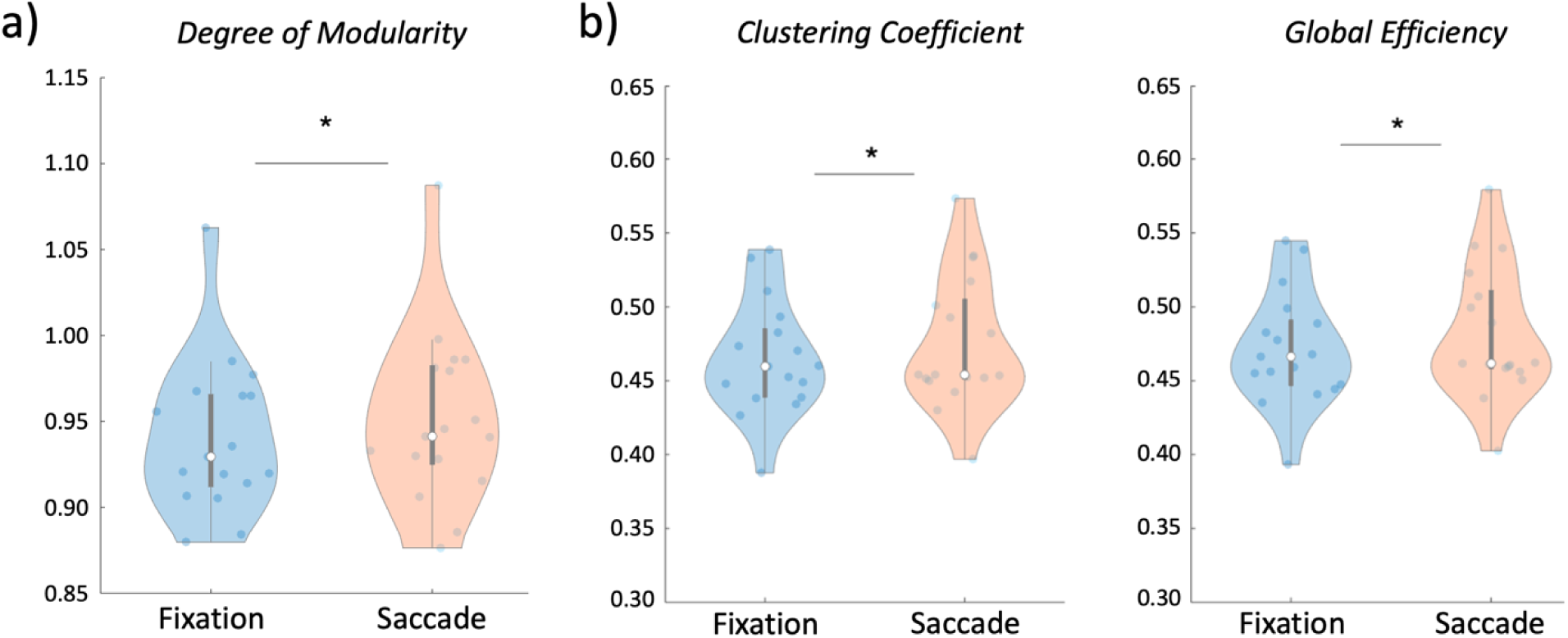
a) Results of the degree of network modularity comparison for the ventral nodes. Conditions were compared using nonparametric permutation t-test (*p* < 0.05). Higher values here indicate greater functional connectivity in the ventral nodes in the *Saccade* condition relative to *Fixation*. **b)** Results of the functional segregation and functional integration measures. Conditions were compared using nonparametric permutation t-tests (*p* < 0.05). *Left*: Clustering coefficient, a measure of overall functional segregation in the network, was greater in the *Saccade* condition relative to *Fixation*. *Right*: Global efficiency, a measure of overall functional integration in the network, was greater in the *Saccade* condition relative to *Fixation*

## DISCUSSION

In this study, we analyzed the network organization of the visual system during an active visual behaviour. Specifically, we subjected correlated BOLD activation of 50 ROIs to a set of GTA measures and contrasted these between our *Fixation* and *Saccade* conditions. Our analysis of the *Fixation* condition revealed three sub-networks: a single bilateral dorsal group, and two lateralized ventral groups (Fig 2) with three important hubs for information propagation throughout the network: bilateral IPS3, IPS4, and V3B (Fig 2). In the *Saccade* condition, the two ventral sub-networks were combined into a single ventral sub-network, consistent with the notion of increased connectivity between the hemispheres for processing information across the two hemifields (Fig 3). Further, we found that the degree of modularity of the ventral nodes increased during saccades, indicating higher functional connectivity (Fig 6a). Additionally, V1v, V1d, V2/V3v, and V2/V3d in the left hemisphere, and V2/V3v in the right hemisphere emerged as important ventral hubs in the saccade condition (Fig 5). Finally, saccades increased overall interconnectivity of the network at both global and local levels (Fig 6b).

Before considering these results further, we acknowledge several caveats and limitations. First, the choice and definition of nodes (Bassett et al., 2011; de Reus & van den Heuvel, 2013; Zalesky et al., 2010). Our selection of 50 nodes was guided by the aim to restrict the data to published, interpretable regions of the visual system, and thus does not represent a ‘whole brain’ analysis (van Wijk et al., 2010; Fornito et al., 2010). Second, this approach and the limitations of our dataset necessitated the use of fixed coordinates, which might obscure intersubject differences (Chong et al., 2017; de Reus & van den Heuvel, 2013; Yu et al., 2018). Third, although we intentionally chose a dataset that incorporates saccades and several object features, our results may not translate to other visual tasks. Finally, GTA essentially relies on signal correlations, and thus cannot prove either anatomic connections or causality. Given these caveats, we will interpret our results with respect to the known physiology and anatomy of system.

### Network Properties in the Fixation Condition

**Dorsal-Ventral Modularity.** The dorsoventral modularity observed here is reminiscent of the two-streams hypothesis of vision, based in the neurophysiology, neuropsychology, and neuroimaging literature (Goodale & Milner, 1992; Goodale & Westwood, 2004; Milner, 2017; Ungerleider & Mishkin, 1982). This hypothesis has been critiqued for various reasons (Pisella et al., 2006; Rossetti et al., 2017; Schenk & McIntosh, 2010), but continues to be an influential principle for shaping our understanding of the visual system.

An advantage of our GTA approach is that it does not rely on (potentially) biased interpretations, but rather on an objective, unsupervised network analysis. Our analysis revealed a dorsal network that spanned nodes and ‘within module hubs’ ranging from parietal areas like V7 and intraparietal cortex to prefrontal areas including the frontal eye fields and dorsolateral prefrontal cortex whereas our ventral modules spanned traditional visual areas (V1, V3, parahippocampal cortex) in occipital-temporal cortex. While this generally supports a two stream hypothesis, it differs from the traditional view of perception-action segregation *within* occipital cortex (Goodale & Milner, 1992; Goodale & Westwood, 2004; Milner, 2017; Ungerleider & Mishkin, 1982). This might be because early dorsoventral modularity requires the presence of an overt action, such as a reach movement (Chen et al. 2014), which did not occur in the current experiment.

Instead, the dorsal-ventral modularity in our task / network may have been driven by the distinction between attention for saccade targets versus visual features (Rolfs and Carrasco 2012; White et al. 2013; Golomb 2019). This notion aligns with the two stream attention system theory (Corebetta & Shulman, 2002), involving a dorsal network for spatial attention, saccade planning / execution, and visual working memory (Jerde et al., 2012; Silver & Kastner, 2009; Vossel et al., 2014) versus a ventral attention system associated with feature processing and (when damaged) hemispatial neglect (Corbetta et al., 2005; Corbetta & Shulman, 2002; Karnath et al., 2001; Mort et al., 2003).

**Lateralization.** Interestingly, our dorsal parietofrontal network was bilateral, whereas our ventral modules were highly lateralized, with the larger ventral module mostly spanning left cortex, and the other smaller module localized to the right hemisphere. This cannot be accounted for by visual field specificity alone, because all three modules showed contralateral specificity at most nodes (Table 1). This seems to suggest that the dorsal module is specialized for spatial computations that inherently require global (cross field) visual processing whereas the ventral networks might be more concerned with local feature processing (Sperry, 1961; Strother et al., 2017; Fabius et al. 2023).

This is again consistent with the notion of a bilateral dorsal attention system that includes IPS and FEF as important regions, and a lateralized ventral attention system (Corbetta et al., 2008; Corbetta & Shulman, 2002; Fox et al., 2006; Vossel et al., 2014). Further, the asymmetric lateralization that we observed in our ventral modules appears to be consistent with the asymmetry observed in hemispatial neglect, i.e., where right damage produces profound localized deficits in the left visual field whereas left damage produces more subtle deficits (Karnath et al., 2001; Mort et al., 2003). Specifically, right hemispheric damage would obliterate our right ventral module, whereas the rightward nodes of the more distributed ‘left’ module would survive left hemisphere damage.

**Global Network Properties and Inter-Modular Hubs.** Finally, our results do not suggest a complete dissociation between modules. Specifically, our analysis of global network properties shows that the entire network of 50 ROIs is both functionally segregated and integrated with heightened interconnectivity at both local and global levels (Fig 6b). This suggests that, despite modular specialization within the system, the overall network remains interconnected to facilitate rapid and effective global transmission of information. This would be expected from a network that must integrate perception and action for real-world behavior (Prime et al., 2011; Rossetti et al., 2017).

More specifically, our betweenness centrality analysis identified bilateral regions of both the dorsal and ventral pathways as significant ‘bridge’ nodes for intermodular communication. These include the intraparietal sulcus (IPS) in the dorsal module and area V3B in the ventral modules (Fig 4). The IPS is considered a significant region of the dorsal attention network that is involved in spatial attention, eye movements, reaching and grasping (Baltaretu et al., 2020; Corbetta et al., 2005; Vesia & Crawford, 2012; Vossel et al., 2014). Conversely, area V3B is known its retinotopic representation of foveal and peripheral vision (Press et al., 2001) and for connecting early visual areas in the occipital lobe with higher order visual processing in the parietal lobe (Arcaro & Kastner, 2015). These regions thus emerge as significant hubs for cross-network communication of spatial attention and working memory (Bressler et al., 2008), and communication between different levels of visual processing.

### Influence of Saccades on the Network

**Modularity**. Our hypothesis was that saccades would increase signal sharing between the two hemispheres, due to the need for integrating visual information from opposing hemifields in the task, a phenomenon referred to as transsaccadic integration (Melcher & Colby, 2008; Prime et al., 2011). We did not observe any major changes in the dorsal network, presumably since it was already bilateral connected. The most dramatic effect of saccades that was the ‘joining’ of the two ventral modules. Transsaccadic feature integration is thought to build on the neural mechanisms for *spatial updating* (Melcher & Colby, 2008; Prime et al., 2007), the process that accounts for changes between the observer and the environment during self-motion (R. F. Wang et al., 2006). This in turn is thought to result in *remapping* of visual signals, which may provide the subjective sense of space constancy during saccades (Sommer & Wurtz, 2008). Previous studies have shown that updating / remapping can occur between the two hemispheres when eye movements displace objects across visual hemifields (Malik et al., 2015; Medendorp et al., 2003; Merriam & Colby, 2005), presumably via the corpus callosum (Berman & Colby, 2009). The observed increase in correlated activity that we observed in our saccade condition, resulting in joining of our left and right ventral modules, is consistent with this idea (Newman, 2006).

**Within and Between-Module Hubs**. Our fixation modules already included hubs that have been implicated in saccades and transsaccadic integration, including the right dorsolateral prefrontal cortex (Tanaka et al., 2014), the frontal eye fields (Ghaderi et al., 2023; Prime et al., 2010, 2011) the lateral and medial intraparietal sulci (Baltaretu et al., 2020, 2023; Dunkley et al., 2016), and dorsomedial occipital cortex (V3B) (Baltaretu et al., 2021, 2023). Most of these hubs persisted during saccades, so to evaluate their specific roles, we contrasted eigenvector centrality between the two conditions.

This contrast identified five major local hubs in the occipital lobe following FDR correction, further highlighting the increased interconnectivity of the ventral nodes, four regions in the left hemisphere (V1v, V2/V3v, V1d, and V2/V3d) and one in the right hemisphere (V2/V3v). This suggests that these areas are more central and interconnected within the ventral modules during saccades. The regions that emerge here are all visual cortex regions that are involved in early processing of visual information for transmission to higher-order processing areas (Goodale & Westwood, 2004). Since this information becomes widely distributed along both dorsal and ventral pathways following early processing in V1, it is reasonable that these early focal points are highly locally connected with those subsequent regions and thus high in Eigenvector Centrality. These visual cortex areas are also heavily involved in remapping during transsaccadic integration, communicating with an intricate network of regions (Malik et al., 2015; Merriam & Colby, 2005). We did not find any significant difference in Betweeness Centrality for Saccades relative to Fixation, but within the saccade dataset we observed that bilateral IPS3 and left V3B’s were important for long-range information propagation across the network.

**Network Properties**. Finally, to analyze the overall local and global properties of the network, *clustering coefficient* and *global efficiency* were used respectively. Clustering coefficient is a measure of functional segregation that provides a value for the degree of interconnectivity at a local scale by investigating closed circuits in the network, with higher values indicating powerful processing at localized levels. Global efficiency, in contrast, is a measure of functional integration that provides a value for the degree of interconnectivity at a global scale by investigating the shortest paths information travels throughout the network, with higher values indicating efficient information propagation between distant nodes. Here, saccadic eye movements led to significant increases in both functional segregation and integration (Fig 6b), indicating that saccades shaped in the network at both local and global levels to maximize efficiency of processing. It is noteworthy that these results are identical to the findings of Ghaderi et al. (2023), despite their use of EEG (as opposed to fMRI) to contrast saccade versus fixation.

These findings are consistent with the idea that transsaccadic integration requires the visual system to integrate multiple information streams (Prime et al., 2008) and again, to share information across visual fields (Medendorp et al., 2003; Merriam & Colby, 2005).

### Conclusions

We employed graph theory analysis to evaluate the modularity and saccade-dependence of fMRI BOLD signals derived from 50 regions of interest, acquired from participants engaged in a shape / orientation change discrimination task: with or without intervening saccades. Our primary results were that 1) in our *Fixation Condition*, three sub-networks of the cortical visual system emerged: a single bilateral dorsal module spanning nodes in parietal-frontal cortex, and two lateralized ventral modules spanning nodes in occipital-temporal cortex, 2) in the *Saccade* condition, the two ventral sub-networks amalgamated into a single, bilateral module, and 3) saccades also increased the overall interconnectivity of the network at both global and local levels. These results provide objective confirmation of signal modularity in the visual system and show that saccades have a profound influence on the network properties in this system.

## MATERIALS AND METHODS

### Participants

On the basis of a power analysis performed with an effect size (0.887) obtained from the most relevant region (right SMG) in a recent transsaccadic study (Baltaretu et al., 2020), we determined that 13 participants would be sufficient to achieve an overall power of 0.914 (Baltaretu et al., 2023). This analysis was performed using G*Power (Faul et al., 2009) with the following parameters: 1) two-tailed repeated measures t-tests, 2) a desired power value of 0.90, and 3) an alpha value of 0.05. 21 right-handed participants with normal / corrected-to-normal vision and no history of neurological disorders were tested. Of these, 17 participants (mean age: 28.7 +/-5.2, 12 females) passed our inclusion criteria for further analysis (Baltaretu et al. 2023). All participants provided informed consent and were compensated for their time. This experiment was approved by the York University Human Participants Review Subcommittee.

### Experimental set-up and stimuli

Participants were required to pass MRI screening and were asked to assume a supine position on the MRI table. Their head lay in a 64-channel head coil. A mount on the head coil was used to support an infrared eye-tracker (for the right eye) to ensure appropriate fixation/eye movements, as well as a mirror that reflected the image of the screen from inside the MRI bore. Participants held a button box with their right hand (index and middle fingers rested on the first and second buttons, respectively). Button press responses were analyzed offline.

The experiment (Fig 1) was conducted in complete darkness. A fixation cross was used and always present (except during the presentation of the mask) to direct participants’ gaze as required by the task. The fixation cross could appear approximately 7° from either the left or right edge of the gray screen (45.1° x 25.1°) along the horizontal meridian. The black stimulus always appeared in the center of the screen and was either a: 1) rectangle, 2) barrel, or 3) hourglass. The dimensions of the rectangle were 12° x 6°; the other two objects had the same area as the rectangle.

### General Procedure

#### Experiment

As show in in Figure 1, an event-related fMRI design was used whereby each trial commenced with a fixation of a cross, presented either to the left or right center, for 7.5 s. Then, the central object (rectangle, barrel-shaped, or hourglass-shaped object) appeared for 3 s at +/-45° from vertical. This was followed by a static noise mask for 0.75 s to avoid an afterimage. The fixation cross would then appear either at the same position as before the mask (Fixation condition) or at the other location (Saccade condition) for 0.75 s. The same object presented at the other possible orientation (Orientation change condition) or one of the remaining two objects presented in the same orientation (Shape change condition) appeared in center for 3 s. The object disappeared and the fixation cross was replaced by an instruction (‘I or O?’) for 3 s, directing participants to indicate if the two objects presented in the trial changed in shape (first button, ‘I’) or in orientation (second button, ‘O’). Thus, there were four main conditions: 1) Fixation, Orientation change, 2) Fixation, Shape change, 3) Saccade, Orientation change, or 4) Saccade, Shape change. These trial types were randomly intermingled within a run (24 trials); there were eight runs in total. Each run began and ended with central fixation for 18 s to establish baseline measures.

#### Imaging parameters

We used a 3T Siemens Magnetom Prisma Fit magnetic resonance imaging (MRI) scanner. Functional experimental data were acquired with an echo-planar imaging (EPI) sequence (repetition time [TR] = 1500 ms; echo time [TE] = 30 ms; flip angle [FA] = 78°; field of view [FOV] = 220 mm x 220 mm, matrix size = 110 x 110 with an in-slice resolution of 2 mm x 2 mm; slice thickness = 2 mm, no gap) for each of the eight runs in an ascending, interleaved manner. A total of 312 volumes of functional data (72 slices) was acquired. In addition to the functional scans, a T1-weighted anatomical reference volume was obtained using an MPRAGE sequence (TR = 2300 ms, TE = 2.26 ms; FA = 8°; FOV = 256 mm x 256 mm; matrix size = 256 x 256; voxel size = 1 x 1 x 1 mm^3^). 192 slices were acquired per volume of anatomical data.

### Analysis

#### Behavioural data

Eye movements and button presses were recorded throughout the experiment, both of which were analyzed offline for correct fixation and production of saccades. Trials where participants made eye movements when not required to were excluded from further analysis. Similarly, trials where incorrect button presses were made were not included in additional analyses. As a result, for 17 participants, 148 trials were removed (out of 3264 trials; 4.5%).

#### Functional imaging data

Functional data from each run for each participant were preprocessed (slice time correction: cubic spline, temporal filtering: <2 cycles/run, and 3D motion correction: trilinear/sinc). A Talairach template (Talairach et al., 1988) was used to transform raw anatomical data. Functional data were coregistered via gradient-based affine alignment (translation, rotation, scale affine transformation). A last step of preprocessing involved applying smoothing using a FWHM of 8 mm.

General linear models (GLMs) were created (BrainVoyager QX 20.6; Brain Innovation) for each run across all participants. Each GLM included four predictors, one for each of the four main conditions: 1) “FixOC”, for fixation trials during which only the orientation of the same object changed; 2) “FixSC”, for fixation trials during which only the shape of the object changed; 3) “SaccOC” for saccade trials where only the orientation of the object changed; and 4) “SaccSC” for saccade trial where only the shape of the object changed. Each of the four predictors were convolved with a hemodynamic response function (standard two-gamma function model) (Friston et al., 1997). Final modifications of GLMs were made to include an “Error” confound predictor for trials during which participants made eye movement and/or button press errors.

#### FMRI signal extraction

To carry out the analyses, spheres were created around a central Talairach coordinate for each *a priori*-defined region-of-interest (ROI; see Table 1). The ROIs were created using BrainVoyager QX v2.8 with a radius of 3 mm. Raw intensity signals were extracted (BrainVoyager QX v2.8; Brain Innovation) from each of these spheres for each run across all participants which were used to carry out functional connectivity analysis.

### Regions of Interest

ROIs were selected based on previously identified regions involved in visual processing and saccades. Several of these regions were identified in the previous study as a result of univariate regional analyses (Baltaretu et al., 2020, 2021, bioRxiv/under review) and we included additional regions based on a probabilistic atlas of standardized visual regions (L. Wang et al., 2015). It was not feasible to reproduce participant-specific localizers for the many visual functions cited here, but we checked which nodes fell within activation produced by the visual field^†^ contrast published in our previous study (Baltaretu et al., 2023). All 50 regions included in the following analyses, their abbreviations, and their Talairach coordinates are summarized in *Table 1*.

**Table 1.**
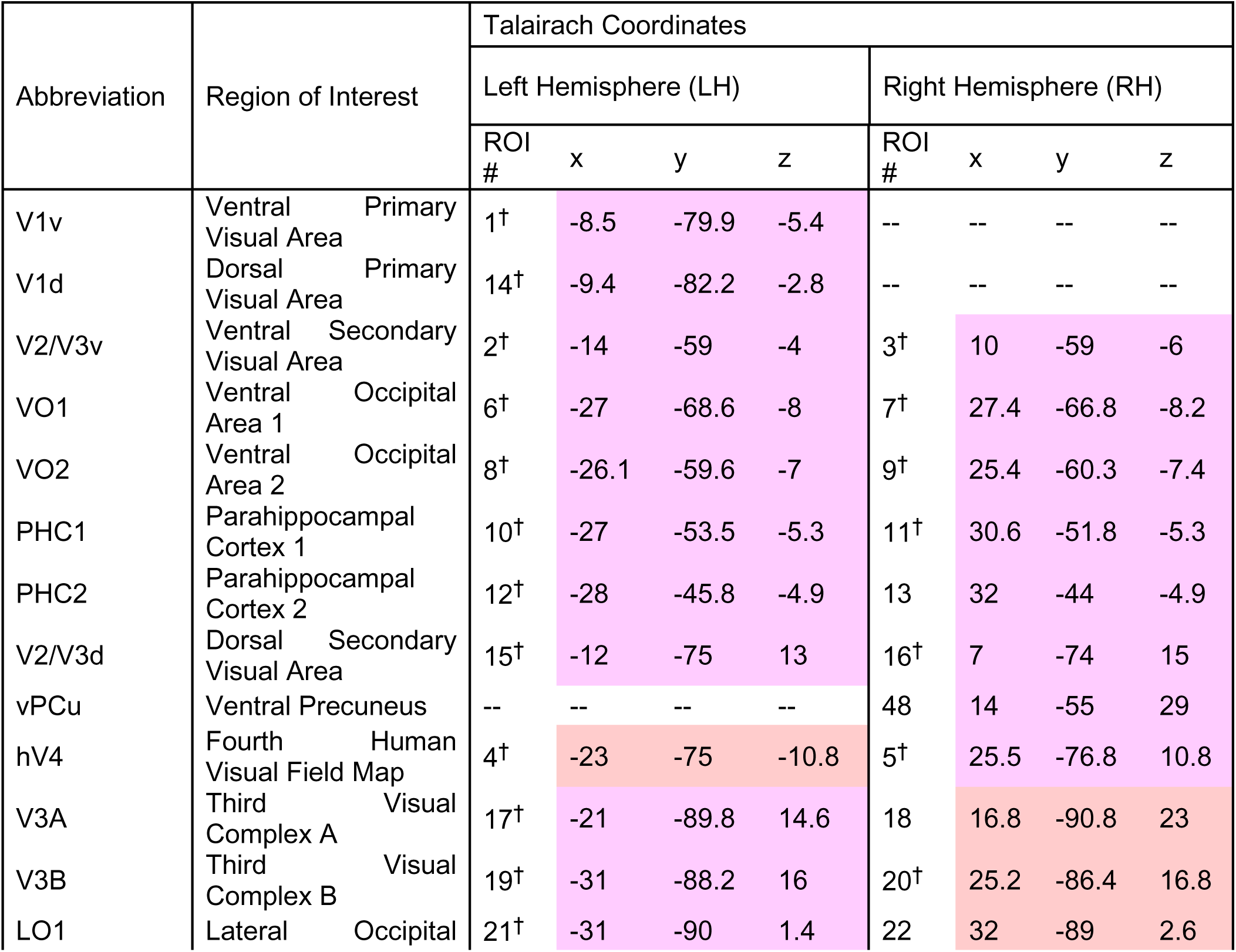
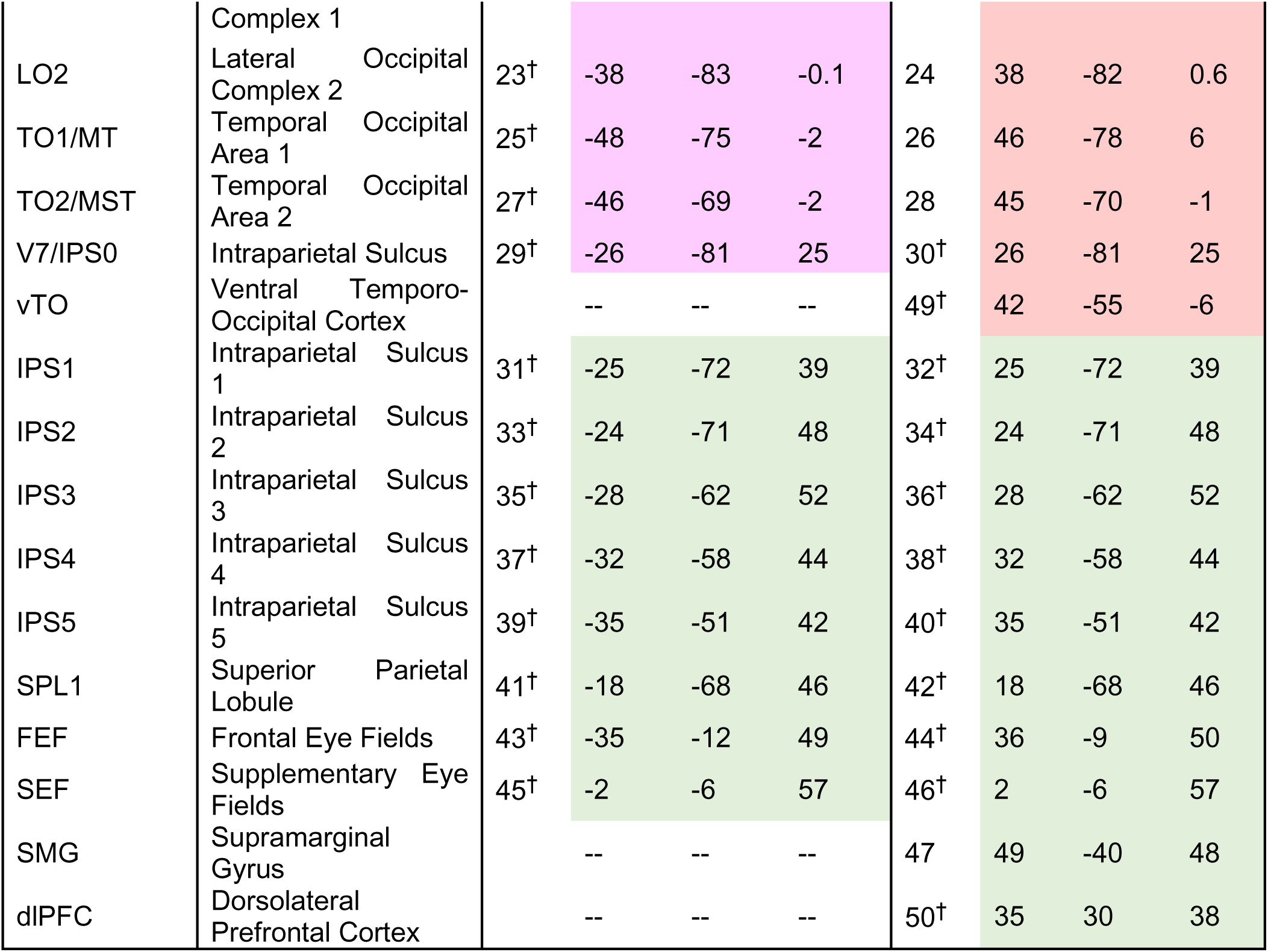
Abbreviations, regions, and Talairach coordinates for each of the 50 ROIs. ROIs are colour coded and organized based on modular affiliation during *Fixation* (mauve: primarily left ventral module, salmon: right ventral module, green: dorsal module) and ‘ROI #’ correspond to the order of rows and columns of adjacency matrices in Figure 7. Superscript symbols^†^ indicate overlap with significant activation produced by visual field contrasts derived from the same dataset (Baltaretu et al., 2023).

### Graph Theory Analysis

To construct the functional networks and compute network properties, we calculated correlations of BOLD activation across our 50 ROIs. Initially, these correlations were conducted for each run and then averaged across runs per *Fixation* and *Saccade* condition for each participant. We then normalized each averaged adjacency matrix by dividing each value by the strongest correlation per participant in each of the two conditions in accordance with best practice guidelines (Rubinov & Sporns, 2011). For each participant, the resulting values were placed in 50x50 adjacency matrices whereby each row (and column) represents a ROI, and their intersection in the matrix communicated the normalized correlation value with each array indicating the functional connectivity between regions; GTA measures were then conducted on these matrices for each participant using the Brain Connectivity Toolbox for MATLAB (Rubinov & Sporns, 2010). Figure 7 shows the weighed, undirected, adjacency matrices of averaged, normalized correlations across participants for each of the *Fixation* and *Saccade* conditions; these averaged matrices were used in the modularity analysis. In this section we present a brief description for GTA.

**Figure 7.**
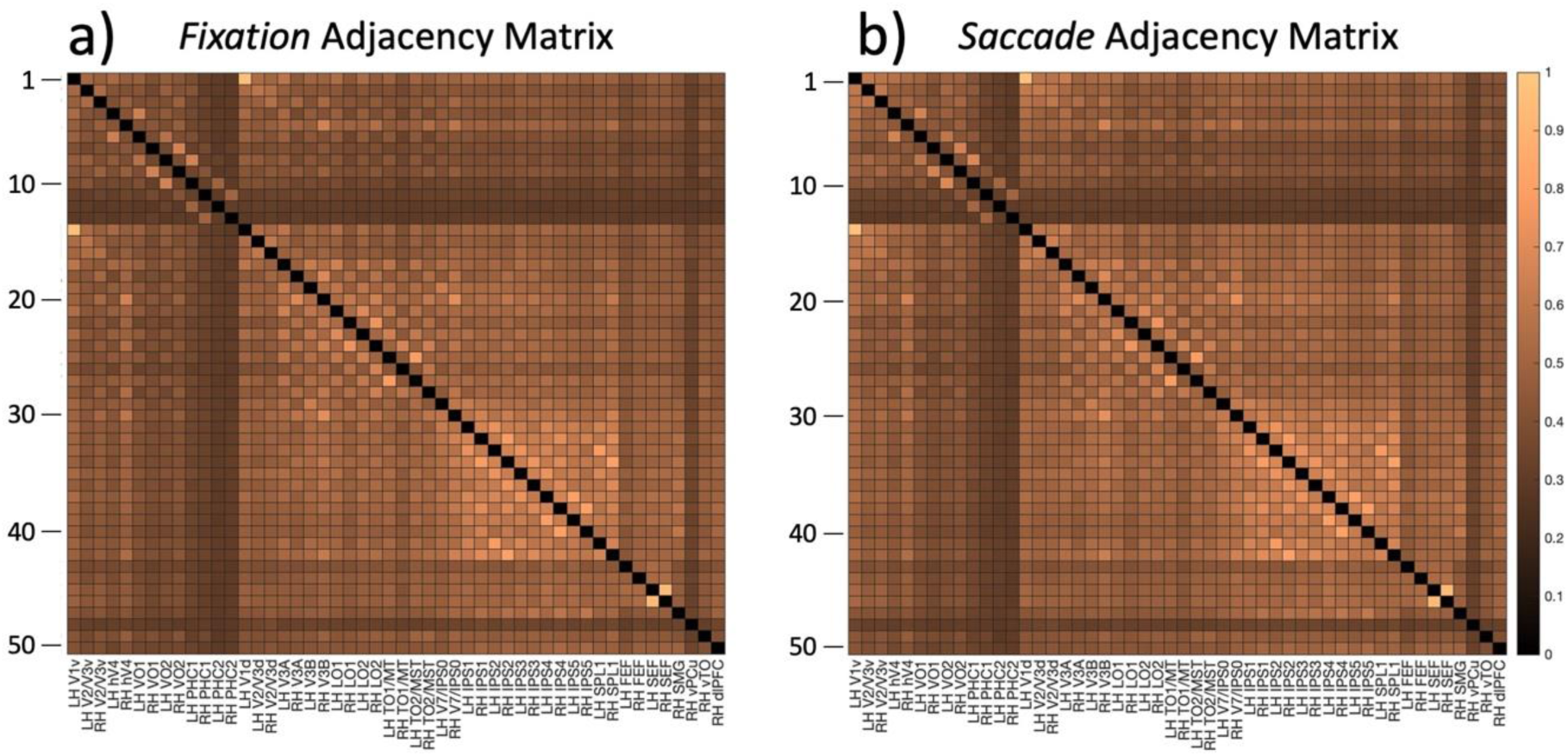
Weighted and undirected adjacency matrices with averaged, normalized correlated BOLD activation across all runs and participants for each of the **a)** *Fixation* and **b)** *Saccade* conditions. Each row (and column) represents a ROI. Intersecting points show normalized correlation value with each region with the other ROIs (*Table 1*), indicating functional connectivity between regions. Brighter cells indicate greater average correlation between regions. The values along the diagonal were set to 0.

*Modularity*, a measure of optimal community structure, attempts to partition nodes in a network into modules consisting of highly interconnected nodes that have few connections with nodes outside of their modular community (Bullmore & Sporns, 2009; Newman, 2004). In the brain, such modules are considered specialized sub-networks of regions organized in such a manner that facilitates information processing among modular regions. Here, we conducted a modularity analysis using the *modularity* function for undirected matrices of the Brain Connectivity Toolbox for MATLAB (Rubinov & Sporns, 2010).

To evaluate modularity of subnetworks, we implemented a Newman modularity algorithm (Newman, 2006) in MATLAB 2019b (open access code for this analysis is available at https://github.com/AHGhaderi/Amir-Hossein-Ghaderi/commit/df636b2105e578a8969ffa88856e54f3267a40a7). The steps to compute modularity are as follows: first, a group of nodes is selected as a subnetwork. For the purposes here, we isolated the nodes from either the dorsal or ventral modules that resulted from the modularity analysis (Fig 2 and 3), and the average connectivity between these selected nodes was calculated. Next, the connections (i.e., ‘edges’) between all nodes in the original network were randomly shuffled and then the average connectivity of our selected dorsal and ventral subnetworks was calculated. With these 2 subnetwork averaged connectivity values calculated, we divided the vales of the original subnetworks and the randomized subnetworks. Significant differences between our subnetworks and randomized subnetwork edges demonstrate that the original subnetworks were modular to a high degree.

*Clustering coefficient* measures functional segregation by quantifying the extent to which a given node’s neighbours also neighbour each other (Bullmore & Sporns, 2009; Watts & Strogatz, 1998), generating closed ‘triangle’ circuits in the network. The mean clustering coefficient for the entire network represents the prevalence of clustering around nodes in the network and high clustering coefficients imply high degrees of functional segregation (Rubinov & Sporns, 2010). In brain networks, these segregated ‘triangle’ circuits account for powerful processing at the local level. Here, we measured clustering coefficient using the adjacency matrices for each condition (*Fixation* and *Saccade*) per participant. Differences in mean clustering coefficient across conditions demonstrate greater interconnectivity between nodes on a local level (i.e., functional segregation) in the condition with greater functional segregation.

*Global efficiency* measures the average inverse shortest path length - the minimal path of nodes and edges that connect two nodes in a network (Boccaletti et al., 2006) - to determine the connectivity between disconnected or distant nodes within a network (Latora & Marchiori, 2001; Rubinov & Sporns, 2010; Watts & Strogatz, 1998). In biological networks, average global efficiency is the network’s ability to transfer information between regions or circuits that are anatomically disconnected but whose communication is necessary to carry out a task. Higher global efficiency is indicative of functional integration of such distant regions (nodes) or circuits of regions (modules). By calculating the difference between global efficiency in our conditions, we can determine changes in functional integration during saccades.

Segregating a network into modules with few connections between sub-networks implies the existence of important ‘bridge’ nodes that serve as connections. We identified these by calculating *Betweenness Centrality*, which calculates the number of shortest paths – paths that connect any 2 nodes with the least number of edges – that cross each node. Nodes with high Betweenness Centrality are considered important for the overall communication of a graph because of the high degree of traffic that passes through them. In the brain, these are important hub regions that relay information and facilitate functional integration between different areas of the brain (Avena-Koenigsberger et al., 2017; Bullmore & Sporns, 2009). Likewise, *Eigenvector Centrality* is indicative of a node’s centrality within a module or cluster of nodes. The degree to which a node is central for the network can vary across different conditions, indicating that different regions are differentially involved in task-dependent information transfer.

### Statistical Analyses

Nonparametric permutation t-tests with 25,000 random iterations were conducted to compare graph theory indices between the *Fixation* and *Saccade* conditions. Nonparametric permutation tests are robust to multiple comparisons and produces results similar to general linear models with multiple comparison corrections (Nichols & Holmes, 2001). For comparison of functional segregation and functional integration, clustering coefficient and global efficiency measures were used respectively. Additionally, Betweenness Centrality and Eigenvector Centrality measures were compared across conditions for each of the 50 ROIs (Table 1) to identify significant hub region changes. To control for these multiple comparisons, false detection rate (FDR) was applied for error correction and an adjusted *p*-value (*padj*) of 0.05 using FDR was adopted for significance (Benjamini & Yekutieli, 2001, 2005; Groppe et al., 2011).

## Acknowledgements

The authors thank S. Sun, Xiaogang Yan, and Diana Gorbet for technical assistance.

